# Breaking Pore Symmetry to Prolong DNA Residence in Solid-State Nanopores

**DOI:** 10.64898/2026.09.20.753066

**Authors:** Simran Nasa, Pranjal Sur, Prabal K. Maiti, Manoj M. Varma

**Affiliations:** Centre for Nano Science and Engineering, Indian Institute of Science, Bengaluru 560012, India; Department of Physics, Indian Institute of Science, Bengaluru 560012, India

**Keywords:** solid-state nanopores, DNA translocation, tilted nanopores, buried membranes, single-molecule sensing

## Abstract

Rapid DNA transport limits the temporal resolution of solid-state nanopore sensing. Here we show that breaking pore symmetry through geometric inclination produces exceptionally prolonged DNA residence without surface functionalization or external trapping. For 500-base-pair double-stranded (ds) DNA, 13 nm diameter SiNx pores in tilted membranes exhibit a distinct long-lived population with a mean dwell time of 11.1 seconds at 600 mV bias, exceeding previously reported dsDNA dwell times by at-least 4 orders of magnitude. Controlled pore tilting reproduces the population of prolonged events, whose prevalence and duration depend on ionic strength. TEM-EELS based thickness measurements exclude substantial membrane-thickness differences, while continuum and atomistic simulations support a mechanism coupling electric-field redistribution to off-axis DNA motion, increased pore-wall contact and suppressed axial mobility. These findings establish geometric inclination as a passive design variable for engineering molecular residence in solid-state nanopores and provide a route towards longer observation windows for single-molecule sensing.

## Introduction

Nanopores provide a direct electrical route for analyzing individual biomolecules by monitoring ion-current changes as DNA, RNA, proteins, or other nanoscale objects pass through a nanometer-scale aperture [1–4]. Biological nanopores have achieved high signal fidelity through their atomically precise channels and well-defined molecular interactions that enable millisecond-scale residence times, facilitating detailed kinetic and structural measurements at the single-molecule level [5]. Despite their advantages, biological nanopores face several intrinsic constraints. Their use of fragile lipid membranes restricts operational stability, especially under high-salt or high-electric field conditions [6, 7]. In contrast, solid-state nanopores offer mechanical robustness, chemical stability, material flexibility, wide pore-size tunability and compatibility with scalable micro- and nanofabrication. [2, 6–9] These advantages make solid-state nanopores attractive for next-generation integrated sensing platforms. However, solid-state nanopores are limited by the exceptionally fast translocation of DNA molecules, many of which typically remain in the pore for only a few microseconds [9]. Together with the higher ionic noise compared to biological systems, this rapid passage restricts the temporal resolution required for extracting fine molecular details [6].

Extensive efforts have been devoted to reducing DNA velocity in solid-state nanopores using strategies that include optical and magnetic manipulation [10, 11] electrolyte engineering using viscosity [12], salt gradients [13], surface functionalization [14], pressure-voltage trapping [15], electroosmotic trapping [16], entropic confinement [17], filtered or double-pore architectures [18, 19] and plasmonic control [20]. While these approaches have led to advancements in the field, they often introduce instrumentation and integration challenges. A complementary design route is to engineer the pore geometry itself so that the transport landscape is modified by fabrication-defined structures rather than by external control.

In our recent work, we introduced a buried silicon nitride membrane architecture in which the freestanding membrane is recessed within a silicon cavity rather than positioned at the chip surface [21]. Anisotropic KOH etching of Si(100) produces sidewalls that are tilted with respect to the Si surface with a characteristic angle of 54.7 degrees introducing an inclined membrane and pore-access geometry. This geometry leads to the question whether pore or membrane tilt, by itself, can alter the local electrostatic and interaction landscape to slow or alter DNA transport? Here we address this question by combining controlled tilted-pore experiments with continuum and atomistic simulations. The experimental design separates three related but distinct cases, namely conventional flat (un-tilted) membranes, membranes intentionally tilted during TEM drilling to isolate pore or membrane orientation as a control variable, and buried membranes [21] that implement a high-tilt membrane with a recessed geometry. These device geometries are referred here as flat, TEM-tilted and buried membranes.

We show that pore and membrane tilt shifts DNA dwell-time distributions towards longer dwell-times due to slower DNA translocation. The observed slowing was as large as four orders of magnitude, which is the largest reported, to the best of our knowledge Supplementary Information Section S6 provides a detailed benchmarking of our work against previous reports. The large increase in residence time observed here cannot be explained from the modest increase in geometric path length alone, motivating a mechanism based on electric field redistribution and enhanced DNA-pore interactions. Finite-element simulations revealed that pore or membrane tilt breaks the symmetry of the local electric field and modifies the electrophoretic force landscape. Molecular dynamics simulations further show that tilted pores favor off-axis DNA configurations, greater contact with the pore wall, and suppressed axial translocation speeds. Thus, we show that pore tilt acts as a passive, geometry-controlled method for translocation speed control in solid-state nanopores without requiring additional forces or surface modifications.

## Results and Discussion

### Design Rationale and Definition of Tilt

Our hypothesis is that a nanopore whose axis or surrounding membrane is tilted relative to the applied electric-field direction can change the capture and transport landscape experienced by DNA. Throughout this manuscript, *θ* denotes the effective inclination of the pore/membrane geometry relative to the wafer/chip surface (Figure 1). The term “flat membrane” refers to a free-standing SiNx membrane at the chip surface with the pore drilled approximately normal to the membrane plane. The term “TEM-tilted membrane” refers to a conventional flat membrane that was deliberately tilted inside the TEM before pore drilling. The term “buried membrane” refers to a recessed SiNx membrane formed inside an anisotropically etched silicon cavity, where the architecture introduces a membrane tilt due to the 54.7 degrees {111} facet angle [21]. The distinction between the TEM-tilted and buried membrane geometries is important because a buried membrane is not merely a tilted version of a flat membrane. It can also modify access resistance, reservoir geometry, hydrodynamic flow, and molecule-wall interactions near the pore. The TEM-tilted membranes therefore serve as a control that tests whether geometric inclination of the pore alone can produce a residence-time increase.

**Figure 1.**
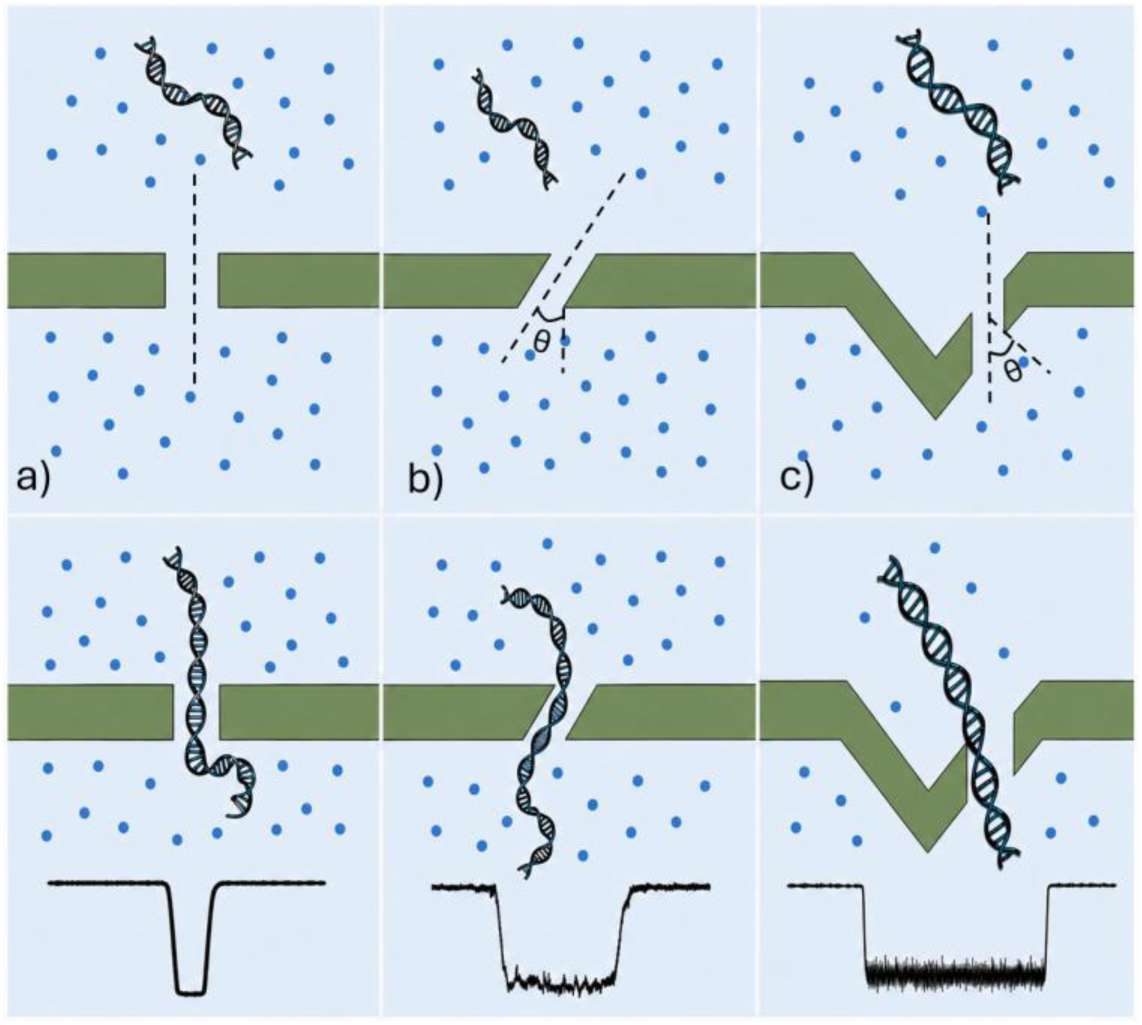
Conceptual basis of tilt-induced residence-time enhancement. (a) In an un-tilted solid-state nanopore (*θ* = 0), the pore axis is aligned with the applied electric-field direction and DNA translocates along a nearly symmetric transport pathway. In a tilted (a) or recessed (c) geometry, the pore-access region breaks axial symmetry, and led to longer dwell times of DNA molecules in the pore during translocations compared to the un-tilted pores.

### Buried Membrane Nanopores Produce Prolonged DNA Residence

We first compared DNA transport through conventional flat membrane and buried membrane nanopores under matched measurement conditions. Nanopores of approximately 13 nm diameter were drilled in both membrane architectures using focused electron-beam exposure in a TEM (SI Figure S1, See SI section S1 for further fabrication details). The membrane thickness in both cases was approximately 20 nm. Translocation measurements were performed with 500 bp DNA in 1 M KCl, 10 mM Tris, and 1 mM EDTA at pH 8.0 at room temperature. Ionic-current traces were acquired at a sampling rate of 200 kHz and low-pass filtered at 10 kHz using an 8-pole Bessel filter.

Flat membrane nanopores produced current blockade events with a median dwell time of around 0.2 ms at a driving voltage of 600 mV. At the same driving voltage, buried membranes also show a cluster of blockade events with a similar median dwell time. However, we also saw a distinct cluster with a median dwell time of 5.2 seconds, which is 4 orders of magnitude higher dwell time compared to the flat membranes (Figure 2).

**Figure 2.**
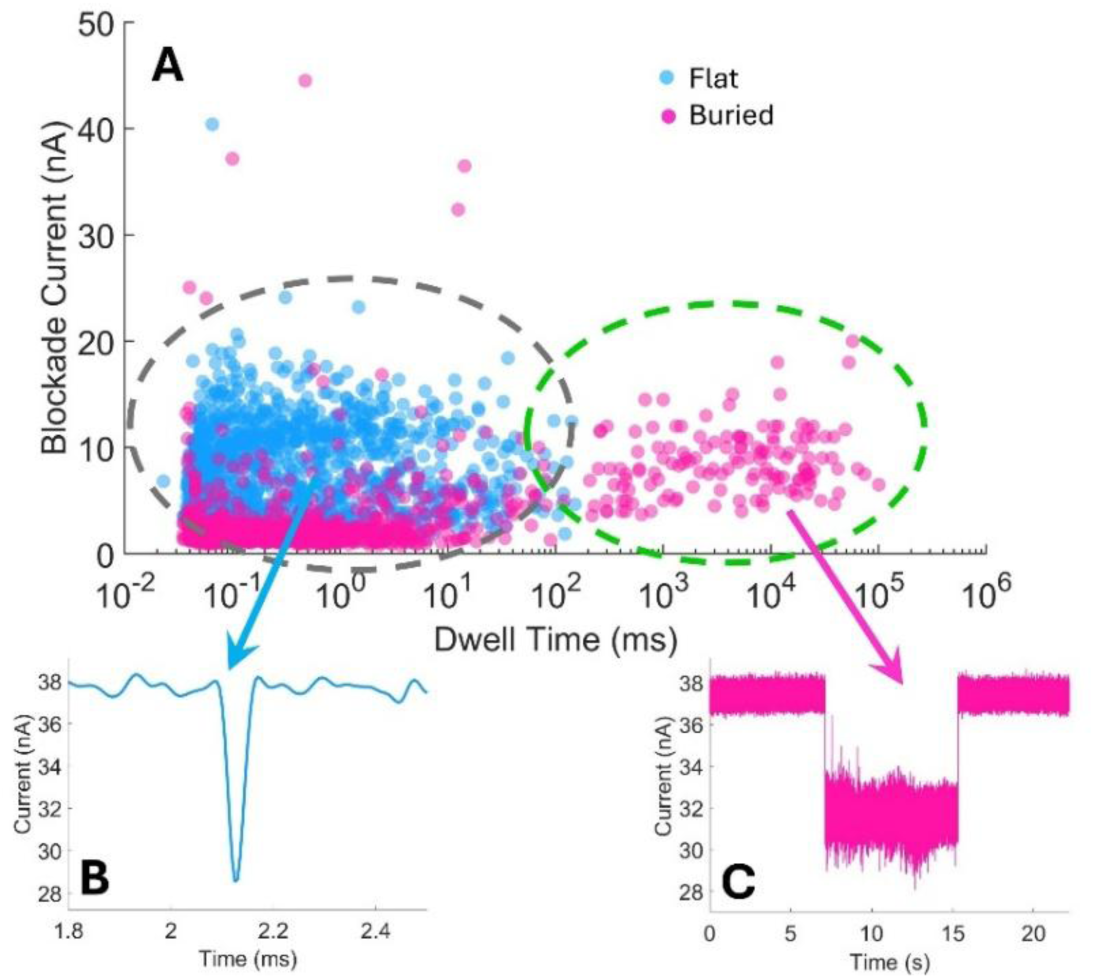
Nanopores fabricated on buried membranes (Figure 1c) exhibit a population of translocation events similar to those in conventional membranes (Figure 1a), as shown by the cluster inside the grey dashed curve. However, in addition to these events, there is also a distinct population of events with 3-4 orders of magnitude higher dwell times (cluster inside the green dashed curve).

For a more quantitative analysis, we modelled the probability distribution function (PDF) of dwell-times from buried nanopores, *f_B_*(*t*) as a linear mixture of the PDF from flat nanopores, *f_F_*(*t*) and a yet to be determined PDF of long events, *f_L_*(*t*). The details of this statistical modelling is provided in SI section S2. One of the challenges of this approach is that as the PDF of flat nanopores, *f_F_* and buried, *f_B_*, are obtained from different chips. As chip-to-chip variation is a confounding variable that could significantly impact the thresholds obtained, an alternate approach is to have a fixed threshold for all data. We used this mixture model analysis to obtain binary classification threshold values of 214.4 ± 44.7 ms for the buried membrane data in the range of 400 to 600 mV (SI Figure S3). Considering this, a fixed threshold of 200 ms was chosen as the criteria to identify events as flat-like, if the detected dwell time was less than 200 ms, or long-events, if the dwell time was more than 200 ms.

### Controlled TEM Tilting Identifies Pore Tilt as a Crucial Variable

To test whether membrane or pore inclination is required to alter DNA transport, we fabricated nanopores in conventional flat SiNx membranes that were intentionally tilted inside the TEM before pore drilling. Membranes of approximately 20 nm thickness were positioned at 0, 10, and 20 degrees relative to the electron beam, and pores with 13 nm diameter were drilled. Because pore formation follows the incident beam direction, tilting the TEM holder during drilling creates a controlled oblique pore geometry as shown in Figure 1b. DNA translocation was analyzed at 400, 500, and 600 mV for flat, TEM-tilted, and buried membrane nanopores under otherwise identical conditions. As shown in Figure 3, flat membranes and 10° TEM-tilted membranes exhibit largely overlapping dwell-time distributions, indicating that small membrane inclinations have negligible influence on DNA transport dynamics. In contrast, nanopores fabricated at 20° tilt (Figure 1b) and buried membrane nanopores (Figure 1c) display broader and substantially extended dwell-time distributions. Increasing the bias voltage from 400 mV to 600 mV increased the number of long-events for the 20° TEM-tilted and buried nanopores. The results shown in Figure 3 demonstrate that membrane or pore tilt systematically slows DNA translocation in solid-state nanopores, with buried membranes behaving similar to pores with higher effective tilt than 20 degrees motivating a mechanistic investigations exploring the origin of these long-dwell time events.

To examine whether electrolyte concentration influences the slowdown observed in buried membrane nanopores, we performed DNA translocation measurements at 0.1 M, 0.5 M and 1 M ionic strengths using 500 bp DNA under otherwise identical experimental conditions. Translocation statistics obtained from flat and buried membrane nanopores were then compared across this ionic-strength range. No measurable translocation events were observed at 0.1 M KCl for either membrane geometry possibly due to reduced blockade current amplitude, reduced translocation time or a combination of both. However, the 0.5 M KCl data exhibited similar behavior as in the 1 M case (Figure 3) with the buried membrane nanopores and 20-degrees tilted nanopores yielding a population of events with substantially longer dwell-times compared to flat membrane and 10-degree tilted nanopores (Figure 4). Importantly, the reduced ionic strength significantly reduced the number of long-events and also shifted the dwell time distributions to smaller values, consistent with previous reports of similar behavior in conventional membranes [22–25].

**Figure 3.**
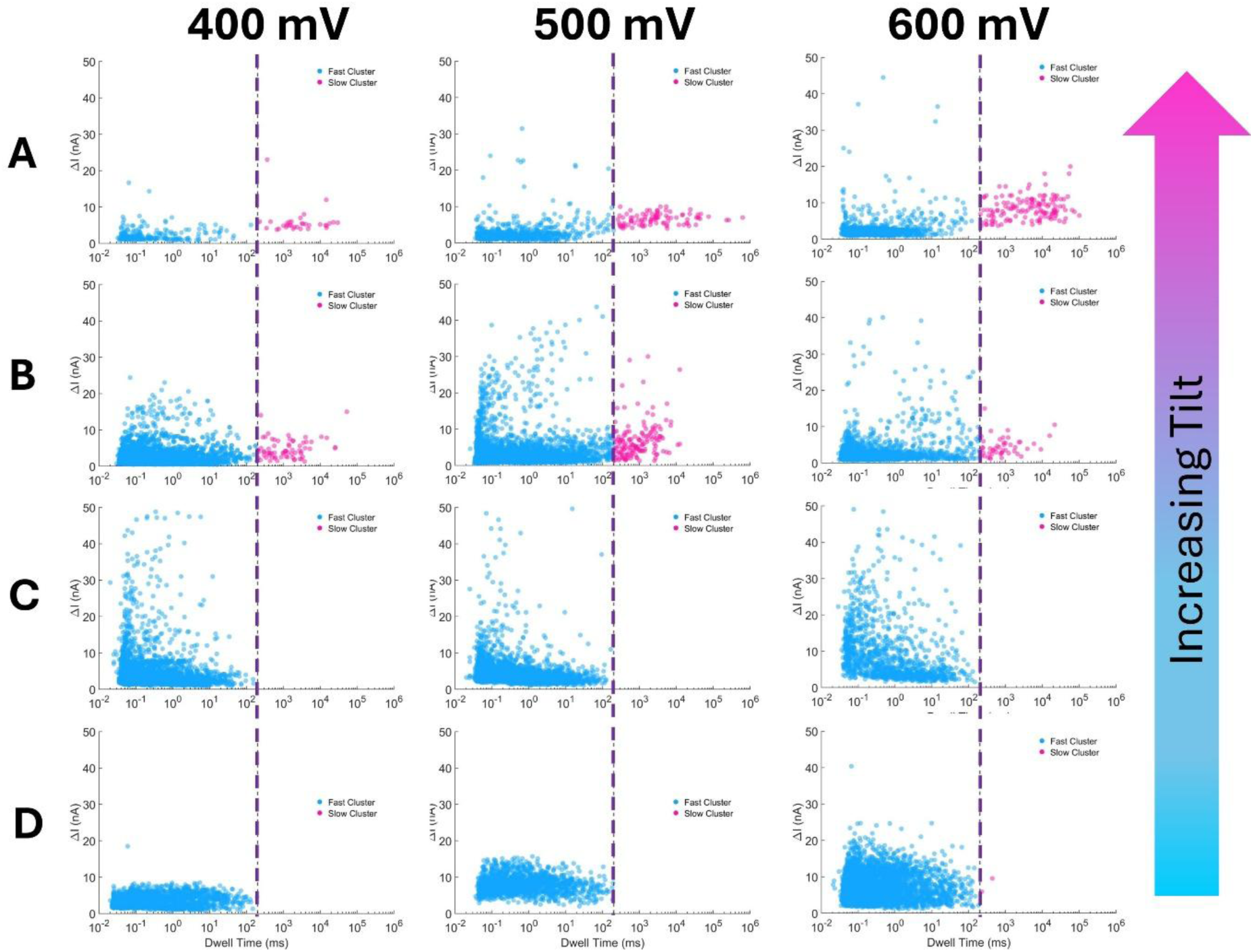
Nanopores on tilted membranes and nanopores tilted related to the membrane produce a population of events with DNA residence times as large as 4 orders of magnitude higher than residence times in conventional nanopores. Buried membranes (Row A) produced a larger number of long-events (residence time > 200 ms) compared to 20° TEM-tilted pores (Row B) while 10° TEM-tilted pores (Row C) and conventional(un-tilted) pores (Row D) do not exhibit this slower translocating population. With increasing bias voltage there was an increase in the number of long-events for the buried nanopores.

**Figure 4.**
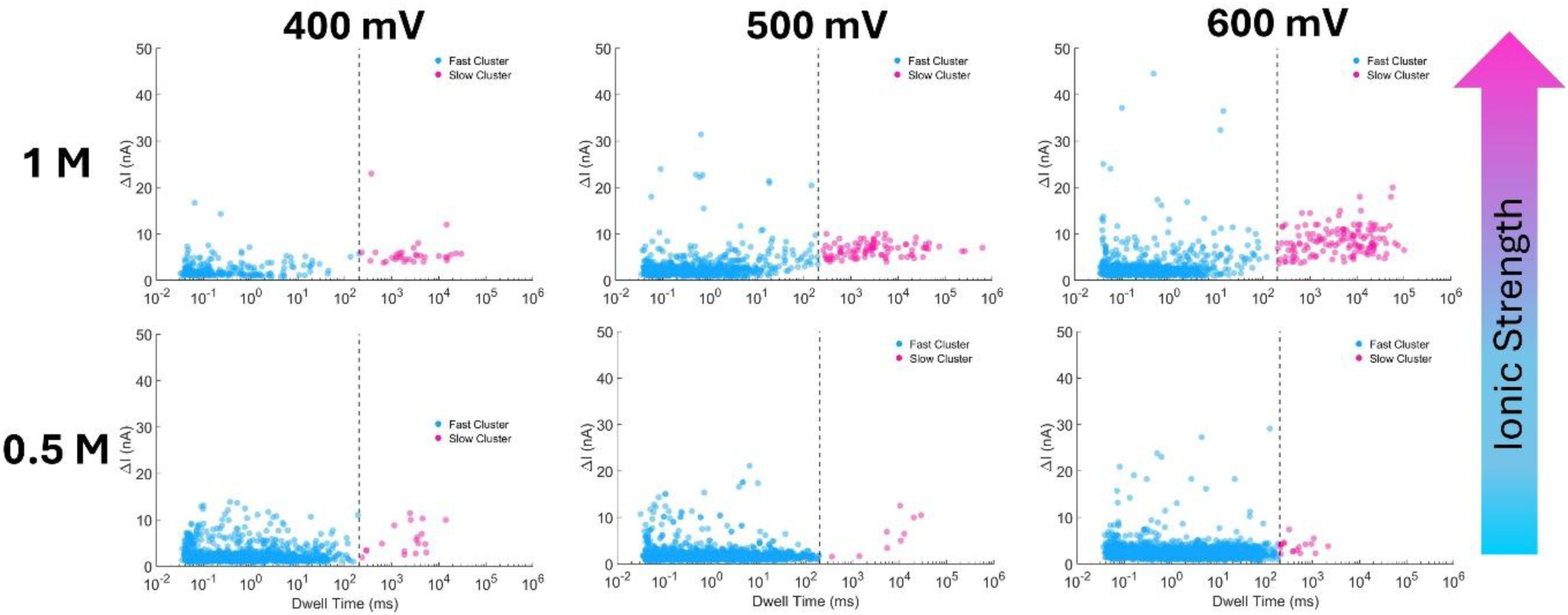
Reducing the ionic strength by a factor of two reduced the fraction of long-dwell time events significantly for the buried membrane nanopores across the 400 – 600 mV bias voltage explored.

Figures 5 and 6 summarise the observations from these experiments using survival probability distributions of the dwell time. The survival probability distribution, *P*(*t*), is simply 1 − *F*(*t*), where *F*(*t*) is the cumulative distribution function constructed from the experimental dwell time data. It represents the probability that a dwell time data point, *t*, will be longer than a specified value *T*, i.e. *P*(*t*) = prob of *t* > *T*. Figure 5 A-C shows that buried membrane nanopores consistently exhibit long dwell-time tails followed by the 20 degrees tilted nanopores while the lower tilt of 10 degrees and un-tilted nanopores do not yield long events across the voltage ranges explored here. Voltage does not appear to have a significant effect on the long events based on the survival probability distributions (Figure 5D-G, and Figure 6G and H) unlike ionic strength where a relatively modest lowering of the ionic strength from 1 M to 0.5 M resulted in a significant decrease in the mean dwell time of the long events (Figure 6 A-F).

**Figure 5.**
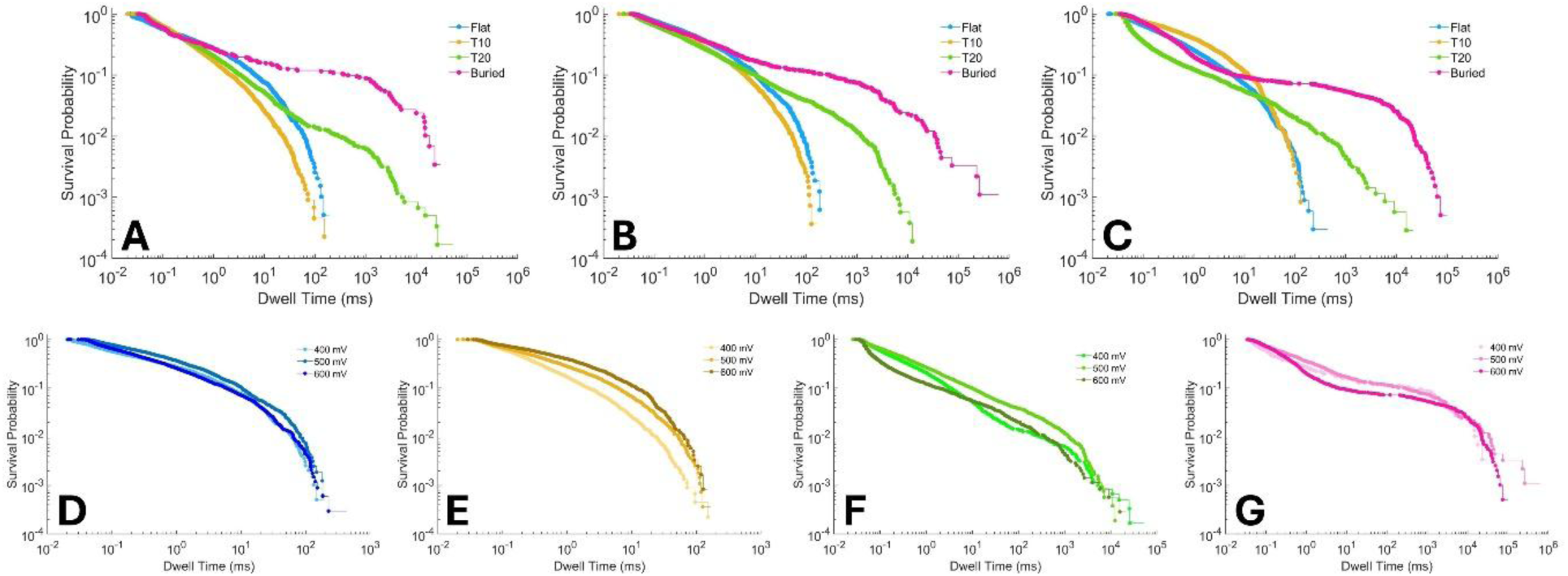
Buried membrane nanopores consistently yielded a larger fraction of long-dwell time events with longer average duration compared to the 20 degree tilted nanopores across the entire bias voltage range explored. Here, A, B, C refer to 400 mV, 500 mV and 600 mV bias voltages, respectively and D, E, F, and G are survival probability distributions for the flat(un-tilted), 10° TEM-tilted, 20° TEM-tilted, and buried membrane nanopores, respectively. The 10 degree tilt was not sufficient to change the translocation dynamics relative to the un-tilted pores for any bias voltage. The electrolyte concentration was 1 M.

**Figure 6.**
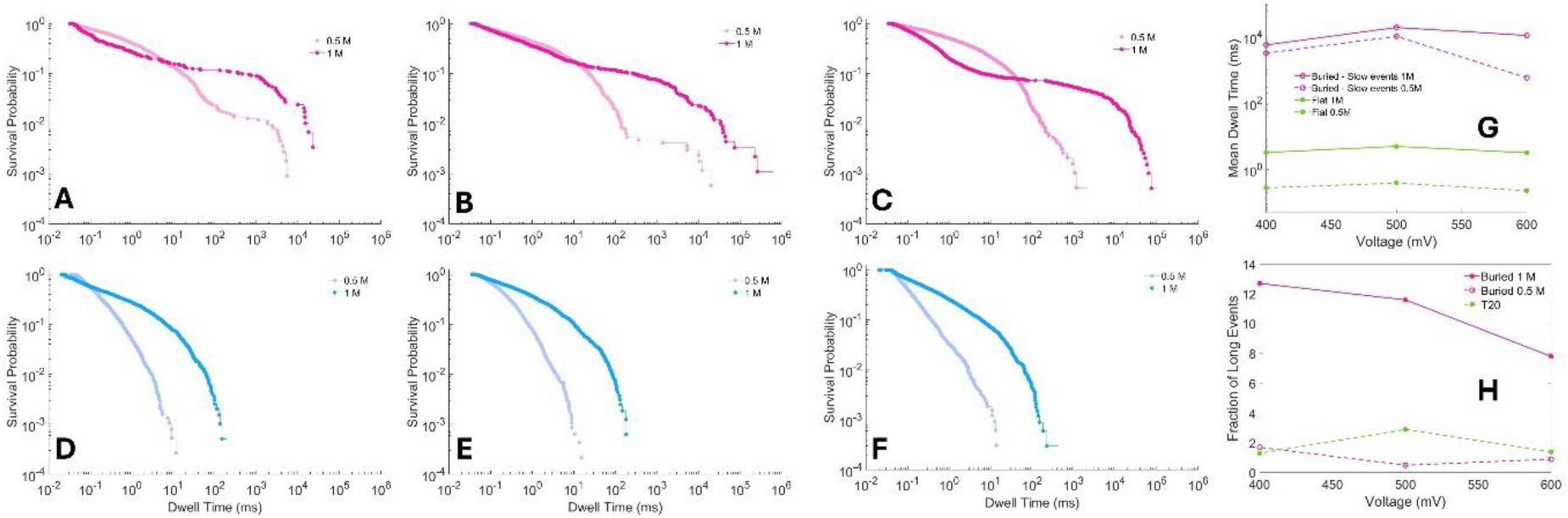
Survival probability distribution of the dwell time in buried membrane nanopores for 400 mV (A), 500 mV (B), and 600 mV (C) bias, indicate that reduction in ionic strength reduced the fraction of long-dwell time events by approximately 6 times and the average duration by a factor of approximately 2 (G, and H). A similar reduction of in dwell time for the conventional un-tilted pores is also visible (D, E, and F representing 400 mV, 500 mV and 600 mV bias voltages, respectively).

Lower ionic strength shifts the dwell time probability distributions to the left, i.e. towards shorter dwell times. This happens for buried as well as flat nanopores. The behavior for the case of flat nanopores has been reported previously and explained based on lower counterion screening in lower ionic strength electrolytes resulting in higher electrophoretic mobility [22–25]. The reduction in dwell time for 0.5 M KCl electrolyte relative to 1 M KCl reported in these previous studies is similar to the 2-3 fold reduction seen in the flat nanopores observed in the present study suggesting a similar mechanism for the effect of ionic strength.

To examine whether the prolonged translocation dynamics observed in buried nanopores could originate from differences in effective membrane thickness, we performed Scanning Transmission Electron Microscopy-Electron Energy-Loss Spectroscopy (STEM-EELS) thickness mapping on both flat and buried membrane geometries [See SI section S3 for further details]. The EELS data did not reveal any substantial difference in thickness between the two architectures (SI Figure S3). These observations suggest that the experimentally observed slowdown cannot be explained solely by variations in membrane thickness, pore path length or other purely geometric aspects of the pore, motivating investigations into the role of electric-field redistribution or other transport variables that may cause the long dwell-time population.

### Continuum Multiphysics Simulation

To elucidate the physical origin of the experimentally observed slowdown, we performed finite-element simulations using COMSOL Multiphysics [See SI section S4 for details]. Stationary continuum fields were obtained by solving the coupled Electrostatics, Transport of Dilute Species, and Creeping Flow modules for two electrolyte reservoirs connected by a nanopore. As simulations using the complete three-dimensional geometries shown in Figure 1, was too computationally expensive, we evaluated the possibility of using an equivalent two-dimensional geometry. We first verified that the electrical potential profiles calculated along the pore axis exhibited qualitatively similar behaviour for the two-dimensional, axisymmetric, and three-dimensional device models (SI Figure S4d). Subsequently, we used the two-dimensional model to do further explorations. Specifically, the simpler geometry allowed us to perform particle-tracing simulations to explore the effect of pore tilt on particle translocation dynamics, which would have been computationally challenging with the full three-dimensional model.

Using the reduced two-dimensional model, we investigated how membrane tilt alters the local electric-field distribution and particle transport dynamics. We observed that, compared to the un-tilted flat pore, increasing the pore tilt causes the electric field to be more confined within the nanopore along with a reduction at the pore entrance and exit (Figure 7B, E, H). The tilted geometry breaks the axial symmetry of the electric field, producing distorted field distributions aligned along the inclined pore orientation. The reduction in the field magnitude at the pore entrance and exit can contribute to prolonged residence within the pore. Equivalent field distortions were also obtained for geometries in which the membrane was tilted relative to a straight pore (SI Figure S5C), indicating that the transport behaviour is governed primarily by the relative pore or membrane orientation.

**Figure 7.**
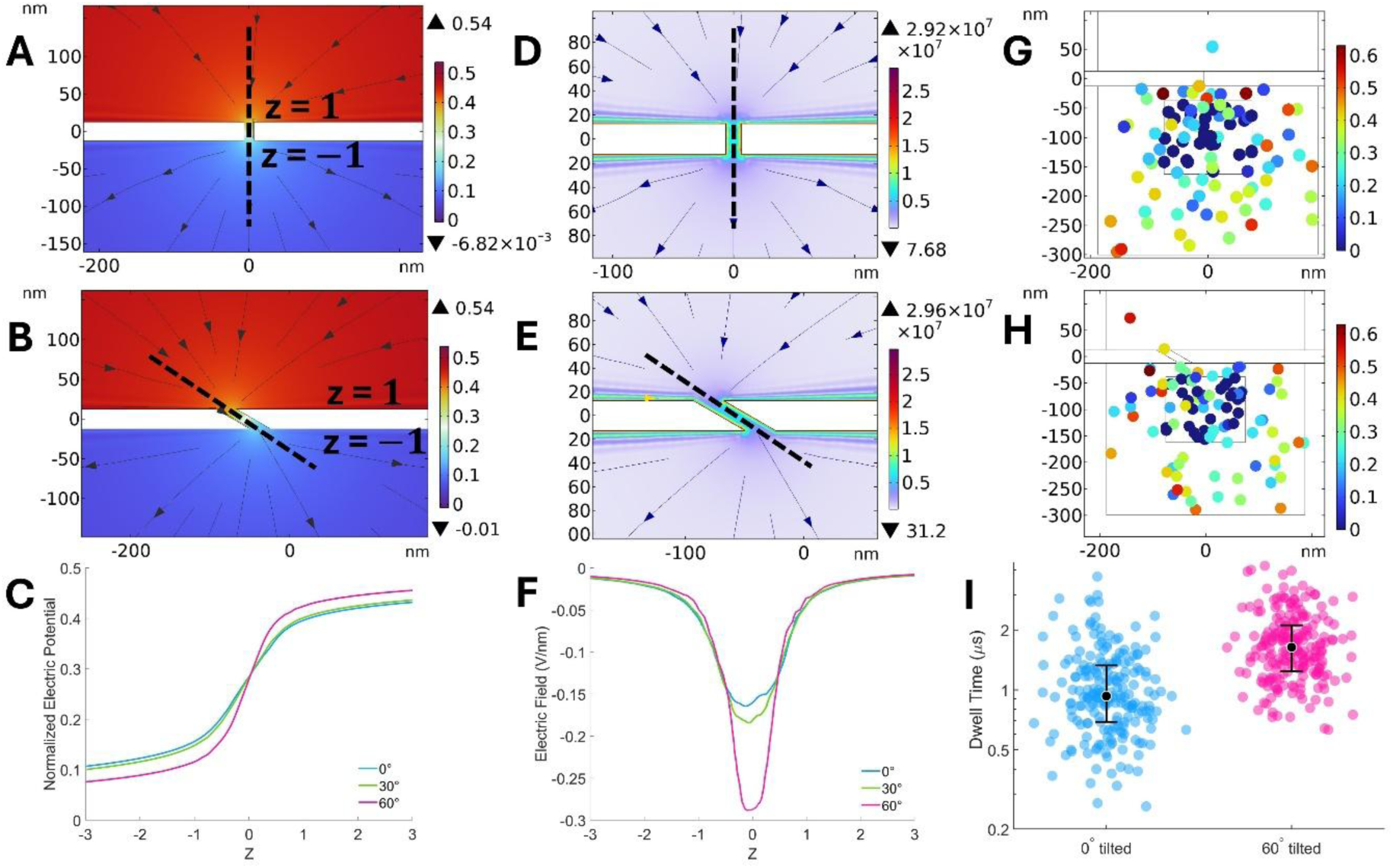
COMSOL simulations illustrating the effect of membrane tilt on electric-field distribution and particle transport dynamics. Electric potential distribution for an un-tilted (A) and tilted nanopore (B) geometries and corresponding electric-field distributions for the un-tilted (C) and tilted nanopore (D) geometries; E and F represent the axial electric potential and electric field profiles extracted from the dashed lines shown in panels A-D. Tilted pore shows a tighter electric field confinement with a reduction in field magnitude near the pore entrance and exit (F); G and H show representative particle configurations during the particle-tracing simulation for the un-tilted geometry and tilted geometries, respectively; the dwell time distribution of the tilted pore is shifted upwards towards longer dwell times compared to the un-tilted pore (I).

To estimate the effect of pore tilt on particle translocation times, the stationary continuum solutions were coupled to particle-tracing simulations in which particles experienced Stokes drag, electrophoretic, and Brownian forces while interacting with pore walls through diffuse scattering [See SI Section 4 for details]. 7C and 7F show representative particle configurations during the early stage of the particle-tracing simulation for the un-tilted geometry for the flat and tilted pores, respectively. We observed that the distribution of translocation times (dwell-times) for the tilted pores shifted towards longer values similar to the experimental observations (Figure 7I). However, the maximum dwell time observed in the case for tilted pore was not substantially larger than the flat pore unlike the experimental observations. This suggests possible role of additional factors such as DNA-pore wall interactions that are not captured in the continuum simulations. To account for these factors, we performed molecular dynamics simulations comparing perpendicular and tilted nanopore geometries.

### Molecular Dynamics Simulations

Molecular dynamics simulations of double-stranded DNA translocating through un-tilted and 30 degrees tilted Silicon Nitride pores were conducted. The simulation geometry is shown in Figure 8A and further details about the simulation methodology are provided in Supplementary Information section S5. The diameter of the pore used in simulations was 4.5 nm and length was 7 nm. Although these dimensions do not exactly match the experimental values of 13 nm and 20 nm, respectively, the smaller dimension of the simulation geometry was needed to keep the computational cost manageable. Further, as the objective of the computational investigations was not to match experimental results with quantitative precision, comparison of the translocation simulations for tilted and un-tilted pores under otherwise identical conditions would reveal qualitative differences and provide insights into the genesis of long-dwell time populations observed with tilted pores. The DNA strand was initially positioned at the pore entrance and application of an electric field along the positive Z axis drove the negatively charged DNA toward the pore interior. Twenty runs for each case, i.e., un-tilted pore and 30 degrees tilted pore, were run with simulation time window of 40 ns each (See SI section S4 for further details).

**Figure 8.**
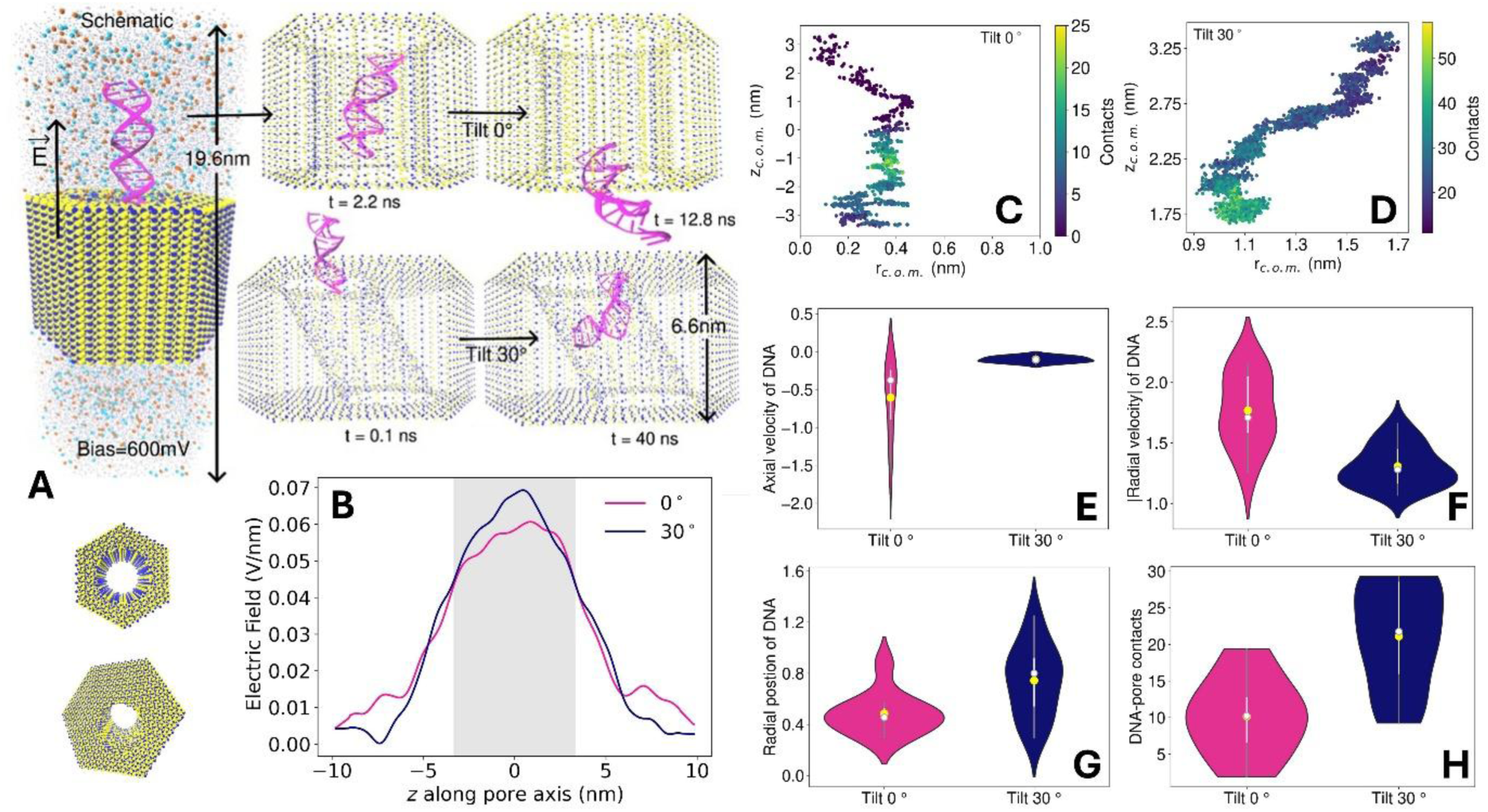
(A) Simulation box used; B) axial electric field profiles show increased field confinement with reduced magnitude near the pore entrance and exit, similar to the multiphysics continuum simulations; C) representative DNA translocation trajectory showing predominantly axial transport in un-tilted pore; D) representative DNA translocation trajectory showing higher radial displacement in tilted pores. This is also true at a statistical level (G); Axial (E) and radial (F) components of DNA translocation are reduced in the tilted pore compared to the un-tilted one; (H) shows increase DNA pore-wall collisions in the tilted pore compared to the un-tilted one.

DNA readily translocated through the no-tilt pore within the simulation time window in 50% of the runs with un-tilted pores whereas we did not observe a single complete translocation of the DNA molecule through the tilted pore [See Supplementary videos SIV1 – SIV8 for representative translocations in un-tilted and tilted pores]. This striking difference is reminiscent of the experimentally observed slow down of DNA through tilted pores. The MD simulations display a tighter confinement of the electric field within the pore and a concomitant reduction near the pore entrance and exit, similar to the continuum simulations (Figure 8B). However, despite the higher electric field inside the tilted pore, the axial and the radial components of the DNA translocation velocity is lower for the tilted pore compared to the un-tilted one (Figure 8E and F). Notably, the axial velocity is suppressed by a nearly an order of magnitude in the tilted pore compared to the un-tilted one. Representative translocation trajectories show a predominantly axial orientation of DNA motion in the un-tilted one (Figure 8C) while, in the tilted pore there is significant motion along the radial direction (Figure 8D), reflecting in an increased average radial displacement of DNA during translocation in the tilted pore compared to the un-tilted one (Figure 8G). The higher radial displacement leads to larger number of DNA pore contacts, indicating the potential for higher DNA-pore wall interactions in the tilted pore compared to the un-tilted pore.

These results indicate that pore tilt promotes DNA conformations and trajectories that favour increased DNA-pore wall interaction, leading to reduced translocation velocity, enhanced dwell time, and suppressed translocation probability.

## Conclusions

This work identifies pore and membrane inclination as a design variable for prolonging DNA residence in solid-state nanopores. Comparisons between conventional, TEM-tilted and buried membrane architectures reveal the emergence of a distinct population of long-duration events, alongside events resembling those in conventional pores. For 500-bp double-stranded DNA, the long-event population in buried membranes reaches a mean dwell time of 11.1 s at 600 mV in 1 M KCl, exceeding previous reported values for dsDNA by more than four orders of magnitude. This enhancement occurs without pore functionalization or auxiliary trapping forces. Ionic-strength dependence indicates that the resulting residence-time distributions are sensitive to the electrolyte environment. Continuum and atomistic computational studies support a mechanism in which geometric asymmetry modifies the local electric field which couples to DNA motion favorinf off-axis configuration that increase DNA interactions with the pore wall and reduce the axial mobility. Although these simulations do not quantitatively reproduce the experimental timescales, they provide a physical basis for understanding how inclination can favour prolonged molecular residence. The primary limitation of the present work is that the prolonged events represent only a minor fraction of the total event population. Also, longer dsDNA fragments, for e.g. 2kb dsDNA, clog the tilted pores inhibiting measurements. Further work is required to engineer the relevant factors such as pore dimensions, alternate symmetry-breaking geometries, and perhaps even surface chemistry to increase the prevalence of the long-lived events observed here. An important next step is also to distinguish extended occupancy in the pore from uniformly slow progression through the pre and examine the impact of this dynamics on the information recoverable from translocating molecules.

## Acknowledgements

We acknowledge funding support from the STARS program of the Ministry of Education (MoE), Government of India under the project STARS-2/2023-0077. We also thank support from MeITY enabling the use of the National Nano Fabrication Facility (NNFC) and Micro and Nano Characterization Facility (MNC), at the Center for Nano Science and Engineering, Indian Institute of Science. We acknowledge assistance from Mr. Anish Kumar Choudhary, master’s student at IIT Gandhinagar as a summer intern in our lab in the curation of dwell-time data reported in literature.

## Data and code availability

The raw data and analysis codes related to this article are available from the corresponding author upon request.

## Supplementary information

### S1. Nanopores fabricated using TEM

Nanopores were fabricated in the SiNₓ membranes by focused electron-beam drilling using a transmission electron microscope (Titan Themis, Thermo Fisher Scientific) operated at an accelerating voltage of 300 kV. The electron beam was focused onto the SiNₓ membrane with an electron dose in the range of 10^8^–10^9^ e^−^/nm^2^ and a full-width at half-maximum (FWHM) of 2–5 nm. Nanopores were initially formed using a tightly converged electron beam, and the pore diameter was subsequently increased by progressively defocusing the beam. The electron-beam conditions were adjusted to obtain a nanopore with a diameter of approximately 13 nm (Fig. S1), which was subsequently used for nanopore characterization and DNA translocation experiments.

**Supplementary Figure S1.**
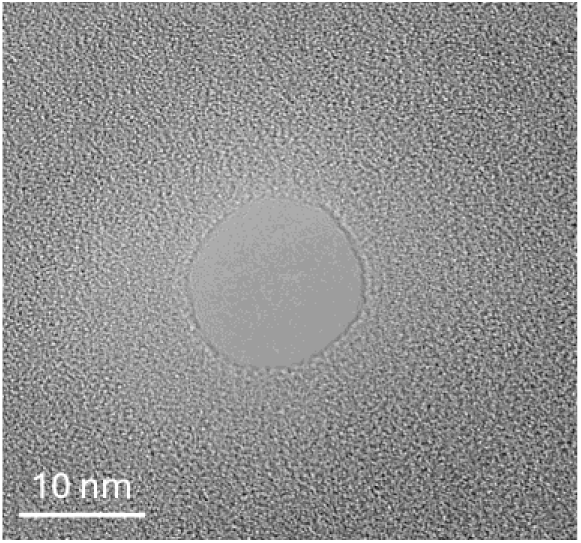
Representative TEM image of a 13 nm diameter nanopore fabricated by focused electron-beam drilling.

### S2. Mixture model to estimate classification threshold

We modelled the probability distribution function (PDF) of dwell-times from buried nanopores, *f_B_*(*t*) as a linear mixture of the PDF from flat nanopores, *f_F_*(*t*) and a yet to be determined PDF of long events, *f_L_*(*t*) as shown in Eq. (S2.1)

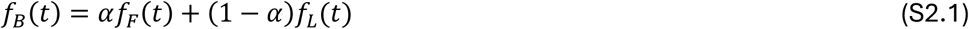

The PDFs *f_F_*(*t*) and *f_B_*(*t*) are approximated by the Kernel Density Estimates (KDE) from the experimentally obtained flat and buried dwell-time distributions. We choose *α* which represents the largest fraction of the buried membrane data that can plausibly be attributed to the flat membrane distribution. Due to the non-negativity of *f_L_* (*t*), the *α* that satisfies this criterion is 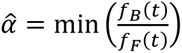. With this choice, the PDF of long-events is obtained as,

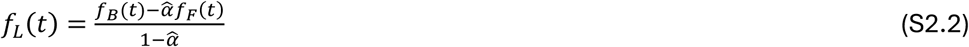

The PDF in Eq. (S2.2) allows us to define a threshold that allows a dwell-time data point to be classified as belonging to a flat-membrane-like population or the long-event population based on the posterior probability 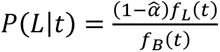. Specifically, the threshold *T* is chosen such that *P*(*L*|*T*) > 0.90 for all *t* > *T*. The exact value of *T* does not change the qualitative conclusions drawn about the flat-like and long-event population because of the persistent criterion that holds for all *t* > *T*. Based on the thresholds obtained for the buried membrane nanopores *T* = 200 ms was chosen as the classification threshold to determine if a dwell-time data point was a flat-pore like event (t < 200 ms) or a long-dwell time event (t > 200 ms).

**Supplementary Figure S2.**
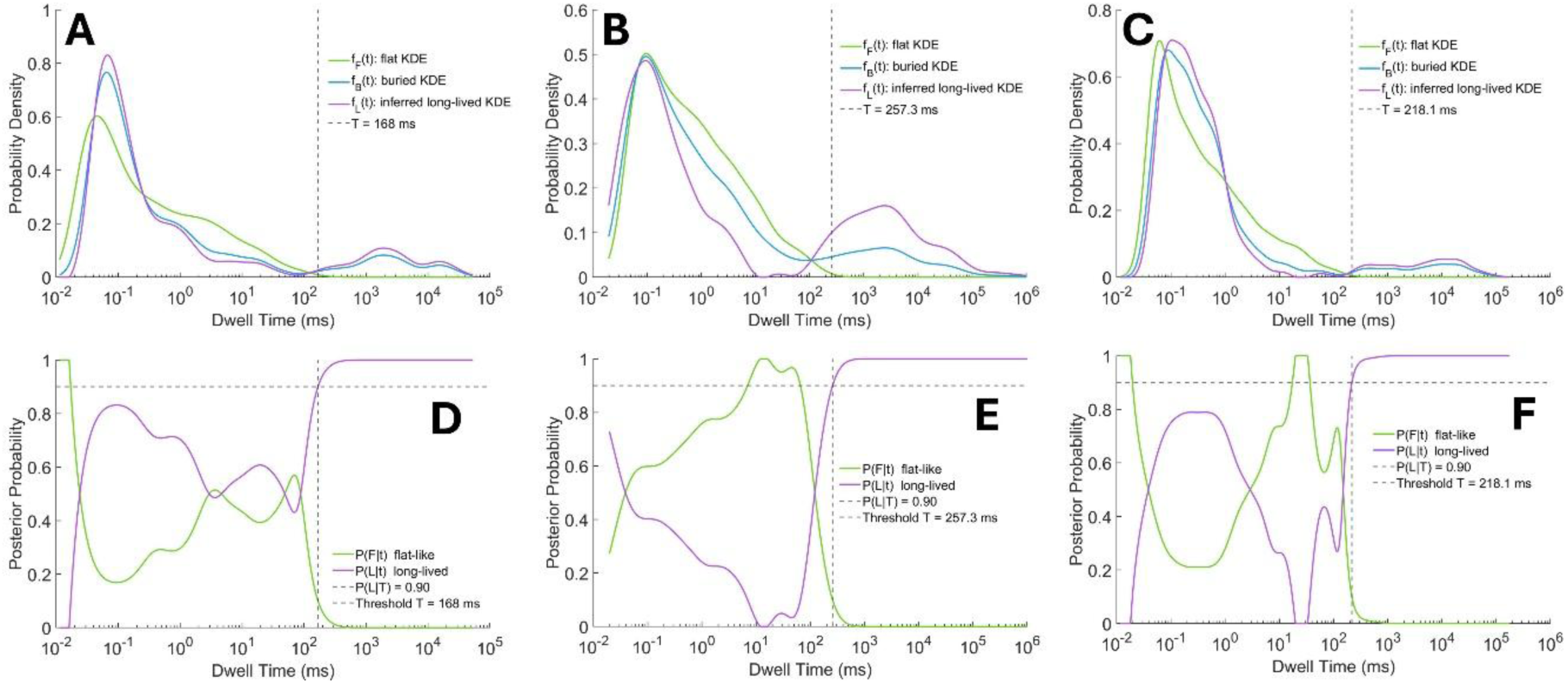
Threshold obtained based on the mixture modeling described above. A, B, and C show the kernel density estimates of the probability distribution functions of the flat (un-tilted), and buried membrane nanopores obtained from experimental data as well as the estimate inferred for the long-dwell time events. A, B, C are for bias voltages of 400, 500 and 600 mV, respectively. Figures D, E, and F represent the posterior probabilities *P*(*F*|*t*) and *P*(*L*|*t*), i.e. the probabilities that a given dwell time comes from a “flat-like” distribution or from the long-dwell time population, respectively. The dashed vertical lines represent the threshold *T*, obtained from *P*(*L*|*T*) > 0.9 for all *t* > *T*. The *T* obtained from these experiments was 214.4 ± 44.7 ms.

### S3. EELS Thickness Mapping of Flat and Buried Membranes

Scanning transmission electron microscopy coupled with electron energy-loss spectroscopy (STEM-EELS) was used to assess whether differences in local membrane thickness could contribute to the distinct DNA translocation dynamics observed in flat and buried nanopores. EELS thickness mapping provides a spatially resolved measure of the relative specimen thickness, expressed as *t*/λ, *t* where is the local specimen thickness and λ is the electron inelastic mean free path. The relative thickness was determined from the low-loss EELS spectra using the log-ratio method

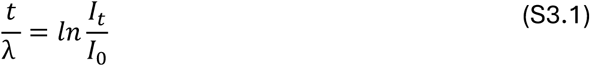

where *I_t_* is the total transmitted electron intensity and *I*_0_ is the zero-loss peak intensity.

STEM-EELS measurements were performed using a Titan Themis microscope equipped with a monochromator, a CEOS probe corrector for Cs-aberration-corrected STEM, and a Gatan Quantum 965 imaging filter for EELS analysis. Low-loss EELS spectral images were acquired over membrane regions containing nanopores with an approximate diameter of 13 nm, with a spatial sampling of approximately 0.8–0.9 nm per pixel and an energy dispersion of 0.1 eV/channel. The resulting *t*/λ maps were used to evaluate spatial variations in thickness across the membrane region, and line profiles were extracted along the indicated directions through the membrane center. The profiles were baseline-normalized to facilitate direct comparison between the flat and buried membrane geometries.

As shown in Figure S3 (a, b) the spatial *t*/λ maps exhibit comparable thickness distributions for the flat and buried membrane regions. The corresponding baseline-normalized line profiles (Figure S3c) show similar thickness profiles across the analyzed regions, with no substantial difference within the experimental resolution. These results indicate that variations in local membrane thickness are unlikely to be the primary origin of the substantially prolonged DNA translocation times observed in buried nanopores.

**Supplementary Figure S3.**
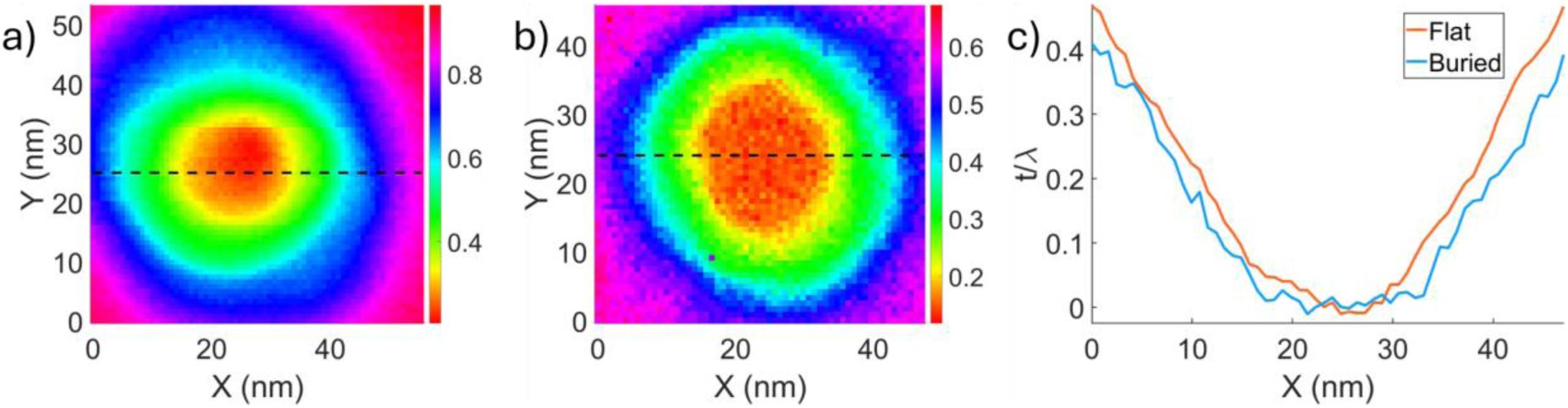
STEM-EELS thickness analysis of flat and buried membrane geometries. (a, b) Spatial t/λ thickness maps obtained from STEM-EELS measurements for flat and buried membrane regions, respectively. The dashed lines indicate the line profiles extracted across the membrane centre. (c) Baseline-normalized t/λ line profiles for flat and buried membranes showing comparable thickness profiles across the analysed regions. The absence of substantial differences in the effective pore-region thickness profile suggests that the experimentally observed slowdown in buried nanopores does not primarily originate from variations in pore geometry alone.

### S4. COMSOL Multiphysics Simulations

#### Simulation Geometry

All simulations were performed in COMSOL Multiphysics version 6.0. The aim was to investigate the influence of membrane tilt on electric-field distribution and particle transport dynamics. The simulation geometry consisted of two electrolyte reservoirs separated by an insulating membrane containing a nanopore (Figure S5a). A reduced-dimensional two-dimensional geometry was employed for the qualitative analysis due to its computational efficiency and consistency with the electric potential profiles obtained from axisymmetric and three-dimensional models (Figure S4). The dashed region shown in Figure S5a indicates the enlarged pore region presented in Figure S5b. To reproduce the experimentally investigated membrane geometries, the nanopore orientation was systematically varied relative to the membrane plane. Simulations were performed for both untilted and tilted nanopore configurations corresponding to the experimentally investigated membrane tilt conditions.

**Supplementary Figure S1.**
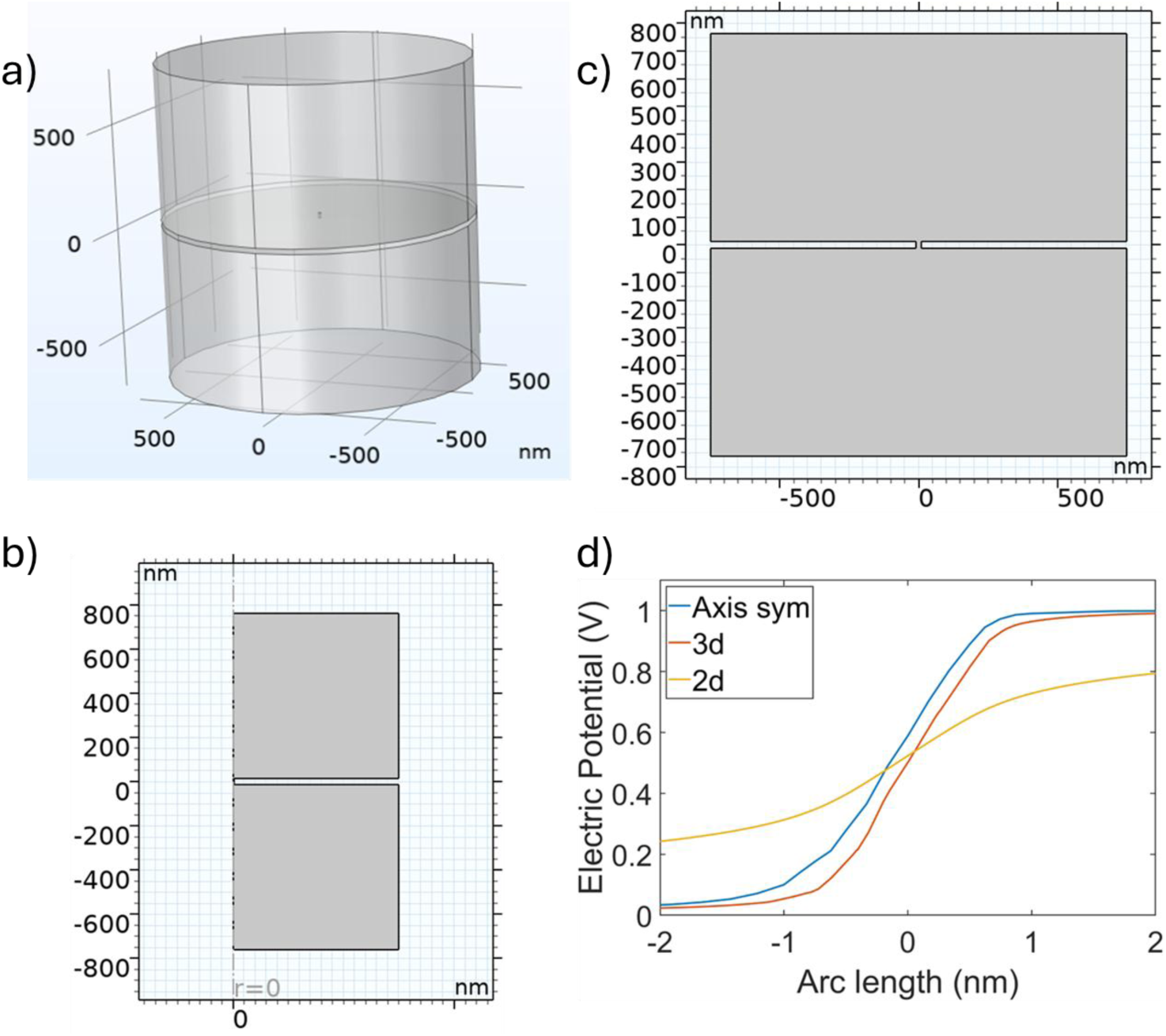
Comparison of COMSOL simulation geometries and corresponding electric potential profiles. (a) Three-dimensional simulation geometry consisting of two electrolyte reservoirs connected by a nanopore. (b) Two-dimensional simulation geometry used for reduced-dimensional analysis. (c) Two-dimensional axisymmetric geometry used for comparison with the full 3D model. (d) Electric potential profiles extracted along the pore axis for axisymmetric, three-dimensional, and two-dimensional geometries, showing qualitatively similar behaviour across all models. The consistency of the calculated potential distributions indicates that the essential electric-field characteristics are preserved in the reduced-dimensional representations used for subsequent particle-tracing simulations

#### Governing Physics and Continuum Fields

Stationary continuum fields were obtained by solving the coupled Electrostatics, Transport of Dilute Species, and Creeping Flow modules. The electrostatic potential distribution generated the electric-field profile used for particle-transport calculations, while the creeping-flow approximation was employed to describe low-Reynolds-number fluid motion within the electrolyte reservoirs and nanopore region.

##### Electrostatics

Electrostatics was used to calculate the electric potential and electric-field distributions governing electrophoretic transport through the nanopore. The electric potential distribution was obtained from Gauss’s law,

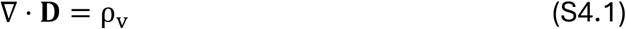

where D is the electric displacement field and ρ_v_ is the free charge density. The displacement field is related to the electric field through D = ɛ**E**, where ɛ is the permittivity of the medium. The electric field field was calculated from the electric potential using

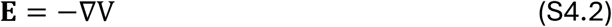

where Vis the electric potential.

##### Transport of Diluted Species

The ionic concentration distribution within the electrolyte reservoirs and nanopore was modeled using the Transport of Dilute Species module under steady-state conditions. Ion transport was described using the steady-state flux equation

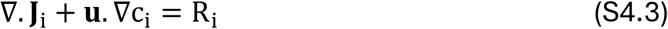

where c_i_ is the concentration of ionic species i, **u** is the local fluid velocity, and R_i_ is the reaction term. The ionic flux was defined as

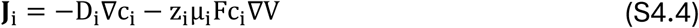

where D_i_ is the diffusion coefficient, z_i_ is the ionic valence, μ_i_ is the ionic mobility, and F is the Faraday constant.

##### Creeping Flow

Hydrodynamic flow within the nanopore system was modeled using the incompressible creeping-flow approximation, appropriate for low Reynolds number transport. Fluid motion was solved using the incompressible Stokes formulation,

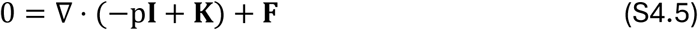

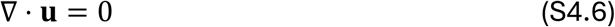

where *p* is the pressure, **I** is the identity tensor, **K** is the viscous stress tensor, **F** is the volume force, and **u** is the fluid velocity.

**Supplementary Figure S2.**
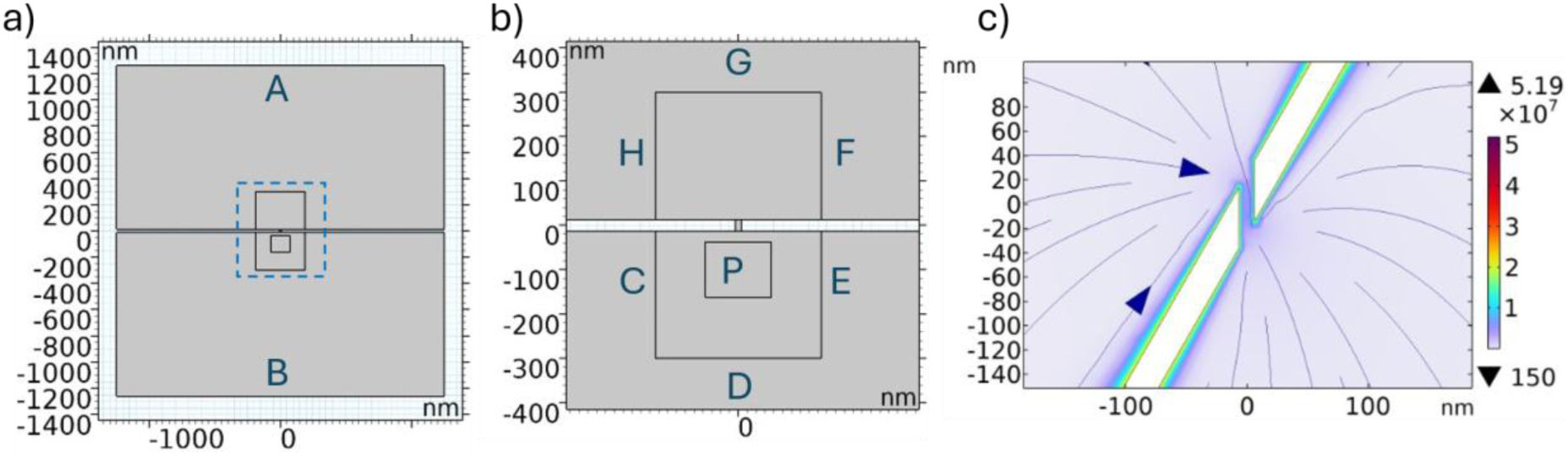
COMSOL simulation geometry and boundary conditions. (a) Two-dimensional simulation geometry consisting of two electrolyte reservoirs separated by an insulating membrane containing a nanopore. The dashed region indicates the enlarged pore region shown in panel (b). (b) Enlarged view of the nanopore region showing the particle-release region (P) and the lateral boundaries (C–H) assigned with disappearing boundary conditions. Boundary A was assigned a fixed electric potential, while boundary B was grounded. (c) Electric-field distribution for a geometry in which the membrane is tilted relative to a straight nanopore axis.

#### Particle-Tracing Simulations

Particle-tracing simulations were performed to qualitatively examine the influence of membrane tilt on transport trajectories and residence times under the combined action of electrophoretic, drag, and Brownian forces. The stationary continuum solution was subsequently coupled to the particle tracing for fluid flow interface.

Spherical particles representing DNA molecules were modeled with a radius of 9 nm and density of 1700 kg/m^3^. Each particle carried a charge of −5*e*, where *e* is the elementary charge. One thousand particles were released from randomized positions within the source region *P* shown in Figure S5b.

Particle motion was governed according to Stokes drag, electrophoretic force, and Brownian force

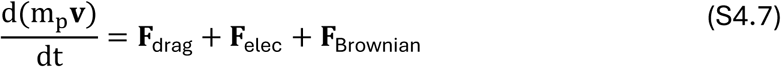

where *m_p_* and **v** are the particle mass and velocity, respectively.

The drag force was implemented using the COMSOL formulationa

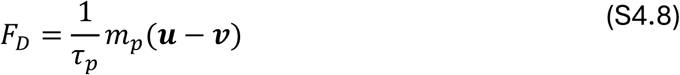

where **u** is the local fluid velocity and τ_p_ is the particle relaxation time given by

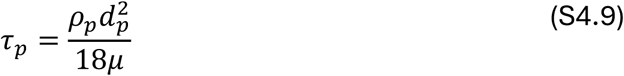

where ρ_p_ is the particle density, d_p_ is the particle diameter and μ is the dynamic viscosity of the fluid. This expression is equivalent to the Stokes drag approximation for low-Reynolds-number particle motion.

The electric force acting on each particle was

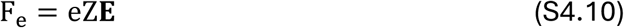

where e is the elementary charge, Z is the charge number of the particle and **E** is the electric field obtained from the electrostatics module. Brownian motion was included through COMSOL’s stochastic thermal force formulation. Particle–wall interactions followed diffuse scattering. Particles entering the opposite reservoir after traversing the nanopore were counted as successful translocation events. The entry and exit times for each particle were recorded, and the dwell time was determined from the time interval between these events. The simulated dwell times increase with pore tilt.

#### Boundary conditions for continuum fields

A fixed electric potential was applied at the inlet reservoir boundary (boundary A in Figure S5a), while the opposite reservoir boundary (boundary B) was grounded. All external lateral boundaries were electrically insulating. To minimize artificial reflections from the finite simulation domain and improve computational efficiency, disappearing boundary conditions were applied at the lateral boundaries labelled C–H in Figure S5b. The membrane and pore walls were treated as solid boundaries and assigned a uniform surface charge density of −23mC/m^2^. No-slip boundary conditions were applied at all solid–fluid interfaces. The electrolyte conductivity corresponded to 1 M KCl. Fluid properties used in the simulations were density 1000kg/m^3^, dynamic viscosity 1.0 × 10^−3^Pa·s, and temperature 300K.

#### Mesh and numerical convergence

The simulation domain was discretized using a physics-controlled triangular mesh with the “extremely fine” mesh setting in COMSOL Multiphysics. Automatic mesh refinement was applied throughout the nanopore and pore-entrance regions to accurately capture steep electric-field gradients and local transport behaviour near the membrane surface. Stationary continuum fields were solved prior to particle-tracing simulations, and the resulting electric-field distributions were subsequently coupled to the particle-transport calculations.

### S5. MD Simulations

#### Simulation procedure

All atom molecular dynamics simulations were performed using GROMACS [1]. The CHARMM27 [2, 3] force field was employed to describe [4–6] the inter-and-intramolecular interactions of DNA. Interaction parameters for Silicon Nitride (Si_3_N_4_) were adopted from the MSXX force-field [27] consistent with previous studies[8, 9]. The CHARMM modified TIP3P water model [10, 11] was used together with the ion parameters developed by Beglov and Roux [12]. Non-bonded interactions were treated using the Lorentz-Berthelot combining rules. The DNA molecule, a poly (ATGC)_4_ sequence was constructed using the Nucleic Acid Builder (NAB) [13] module of AmberTools [14]. Detailed information on the simulation systems and run statistics is provided in Figure S6, and Tables S1 and S2. Periodic Boundary Conditions were applied in all three spatial directions. The Settle algorithm [15] was used to maintain the rigidity of the water molecules, while LINCS [16] was employed to constraint all bonds involving hydrogen atoms.

Energy minimization was carried out using the steepest descent algorithm for 50,000 steps. The minimized system was subsequently equilibrated in the NVT ensemble for 2 ns at 300 K using the Bussi-Donadio-Parinello thermostat [17] with a time constant of 0.1ps. This was followed by NPT equilibration for 2 ns to achieve equilibrium density at 1 bar reference pressure using semi-isotropic pressure coupling with the Parinello-Rahman barostat [18] (time constant = 2ps). During both equilibration phases, the DNA molecule was position restrained with a force constant of 1000 kJ/mol.nm^2^. Production simulations were performed in the NVT ensemble using the Bussi-Donadio-Parinello thermostat with a time constant of 1 ps, maintaining the system temperature at 300 K. The integration time steps were 2 fs during both equilibration and production phases, and trajectories were saved at 10 ps intervals for analysis. Short-range Lennard–Jones and Coulomb interactions were computed using a cutoff distance of 12 Å, while long-range electrostatic interactions were treated using the Particle Mesh Ewald (PME) method [19]. An external electric field of 0.031 V/nm was applied along the +Z direction during the production runs, corresponding to a total bias of 0.6V across 19.6 nm long simulation box. Harmonic restraint of force constants 1 kcal/mol. Å^2^ and 10 kcal/mol. Å^2^ were applied to the bulk and surface atoms of Si_3_N_4_ membrane, respectively throughout all stages of the simulations. The time-dependent ionic current through the nanopore was calculated as [20],

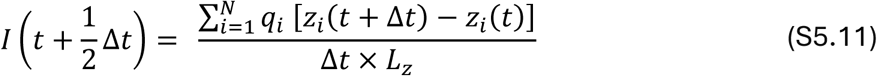

where *I* is the ionic current, Δ*t* is 2 ps, and *z_i_* is the *z* coordinate of the *i^t^*^ℎ^ ion, and *q_i_* is its charge. N is the total number of ions, and the sum is for all the ions. The relative current blockade has been calculated as (*I*_0_ − *I_avg_*)/*I*_0_, where *I*_0_ is the open-pore current. Trajectory visualization was performed using Visual Molecular Dynamics (VMD) [21–23], while trajectory processing and analysis were carried out in MDAnalysis [24, 25] environment. The electrostatic potential maps were computed using the PMEPot plugin in VMD [26], and the corresponding electric field maps were obtained by taking the negative gradient of the electrostatic potential.

**Supplementary Figure S3.**
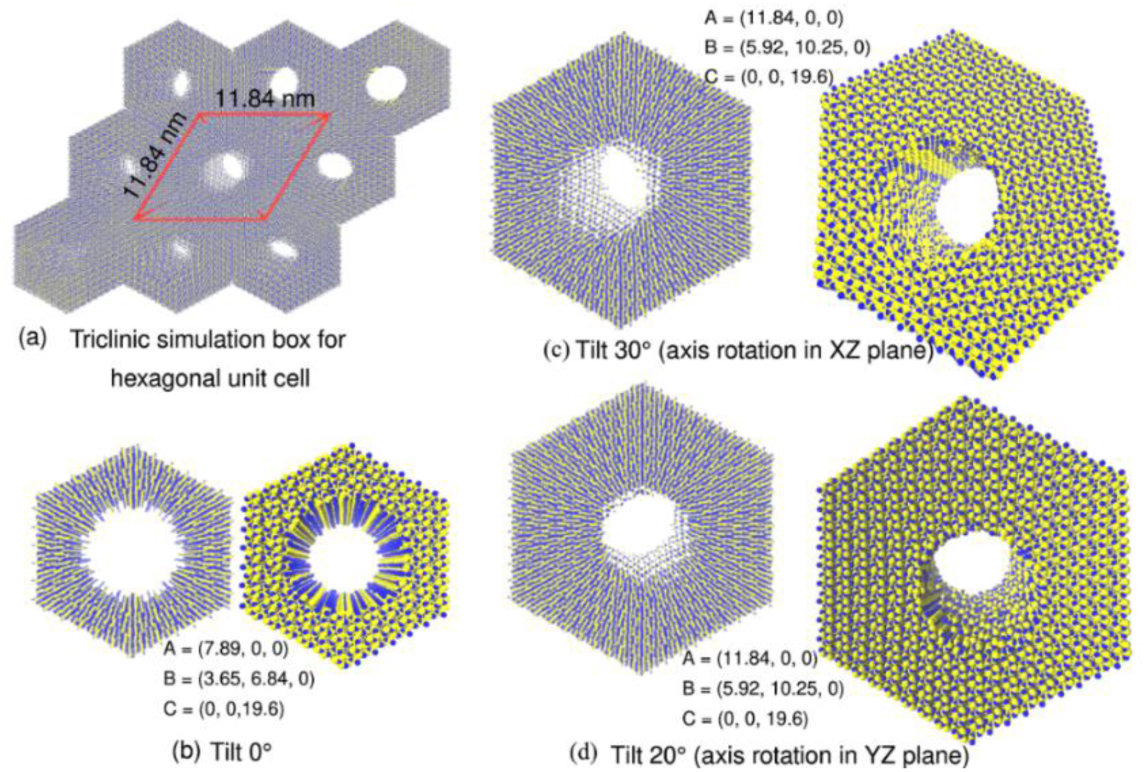
(a) Triclinic periodic boundary condition for SIN. The simulation box dimensions are depicted. All the box vectors are in nm. The pores are shown in different representations and angles in the following: (b) Perpendicular pore, (c) Tilted pore with a 30° tilt and the rotation about Z axis is in XZ plane, (d) Tilted pore with a 20° tilt and the rotation about Z axis is in YZ plane.

**Table S1:** Number of atoms in the simulation system.

DNA Translocation
| Pore | K<sup>+</sup> | Cl<sup>-</sup> | Water | Si<sub>3</sub>N<sub>4</sub> | DNA | Total |
| --- | --- | --- | --- | --- | --- | --- |
| Tilt 0° | 647 | 617 | 74742 | 19280 | 1014 | 96300 |
| Tilt 30° | 1419 | 1389 | 159069 | 55314 | 1014 | 218205 |

Open-pore
| Pore | K<sup>+</sup> | Cl<sup>-</sup> | Water | Si<sub>3</sub>N<sub>4</sub> | Total |
| --- | --- | --- | --- | --- | --- |
| Tilt 0° | 617 | 617 | 75909 | 19280 | 96423 |
| Tilt 30° | 1389 | 1389 | 160188 | 55314 | 218280 |
| Tilt 20° | 1389 | 1389 | 159186 | 56353 | 218317 |

**Table S2:**
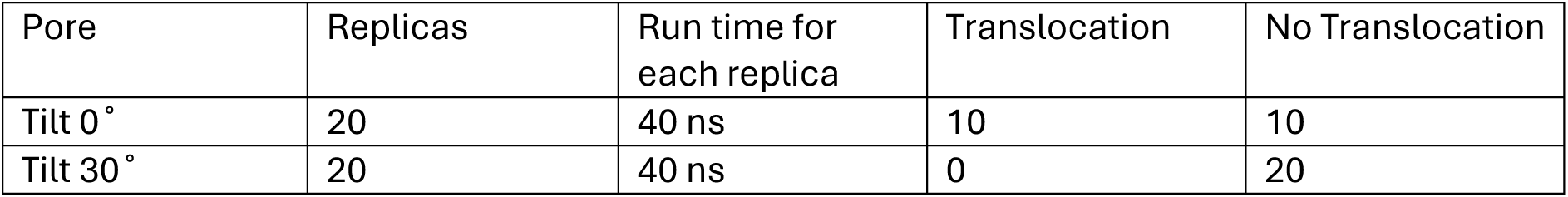
Production-run statistics for translocation studies.

| Pore | Replicas | Run time for each replica | Translocation | No Translocation |
| --- | --- | --- | --- | --- |
| Tilt 0° | 20 | 40 ns | 10 | 10 |
| Tilt 30° | 20 | 40 ns | 0 | 20 |

Here, a translocation is considered successful if the center-of-mass (COM) of the DNA comes out of the pore on the trans side.

**Supplementary Figure S4.**
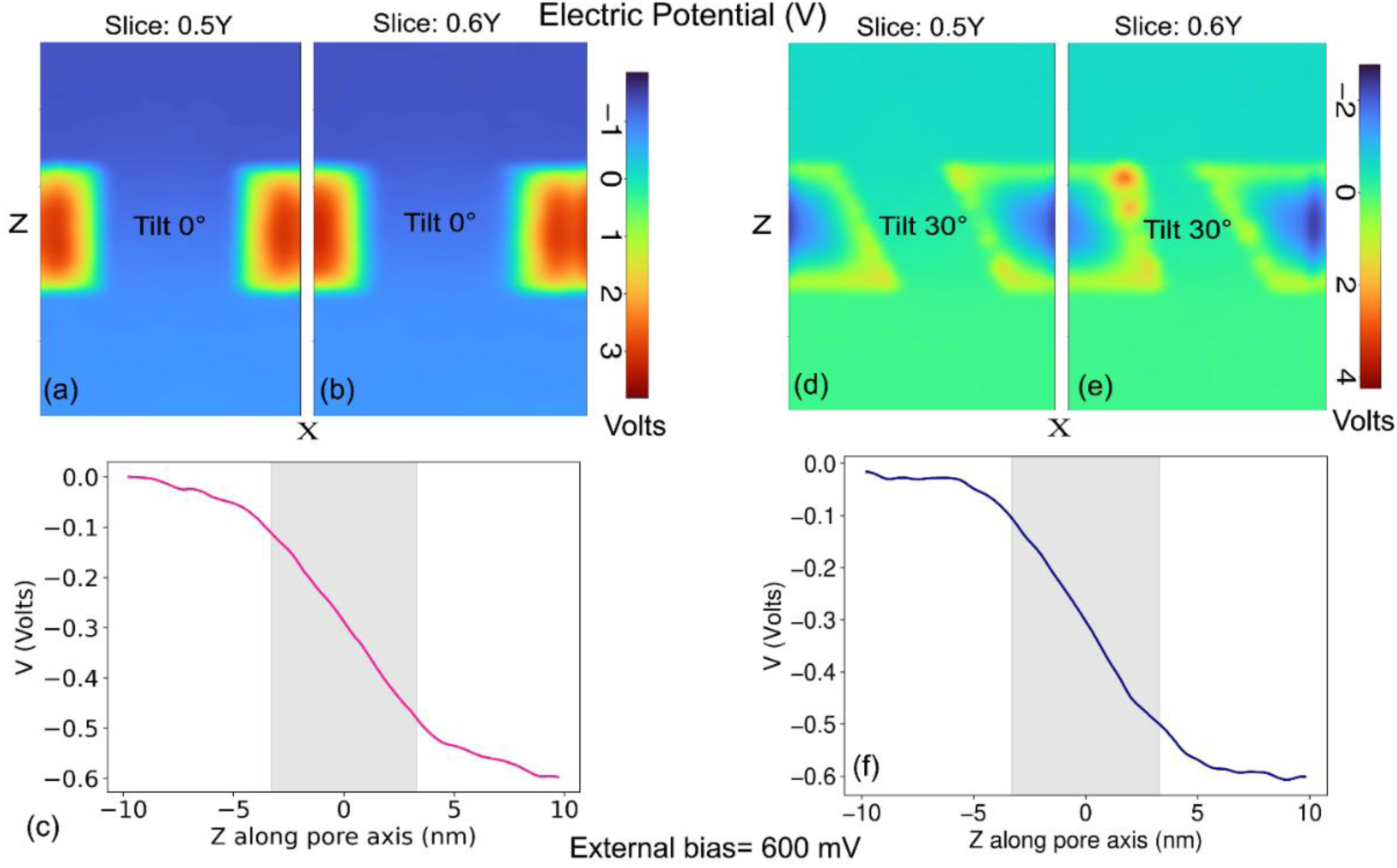
(a)-(b) Electric potential distribution on the middle XZ plane and on the XZ plane at 0.6Y, respectively inside the perpendicular pore. Moving away from central plane does not change the symmetry of distribution much. (c) Average potential profile close to the axial line for the perpendicular pore. (d)-(e) Electric potential distribution on the middle XZ plane and on the XZ plane at 0.6Y, respectively inside the tilted pore. Moving away from central plane increases the asymmetry of distribution. (f) Average potential profile close to the axial line for the tilted pore.

### S6. Dwell-time Benchmarking

A comprehensive literature survey was conducted to compile experimental data on DNA translocation speeds for solid-state nanopores. The search strategy included both backward and forward citations of key studies to ensure extensive coverage of original works and subsequent developments. Primary criteria for inclusion were studies reporting measurable translocation dynamics, specifically, works that actively sought to slow down DNA translocation in solid-state nanopores. For each selected paper, the characteristic dwell time associated with DNA translocation was extracted. Data for systems employing both ssDNA and dsDNA were used. Since individual studies report these times differently, such as, mean translocation duration, exponential decay time, or time constant these data were treated as the representative of the characteristic dwell time. When multiple characteristic times were reported for distinct translocation event types within a single study, the event type exhibiting the highest dwell time was selected as representative. These studies have been done across a range of DNA length, bias voltage and pore diameters. These parameters were also noted, and this dataset was used for benchmarking the long-dwell time observed in this study. As the representative dwell-time value for our study, we used the mean dwell time of the long-dwell time event population for the buried membrane nanopores at 600 mV bias voltage which was 11120 ms. The DOIs of the 37 references used for benchmarking is provided in SI Table S3.

**Supplementary Figure S8.**
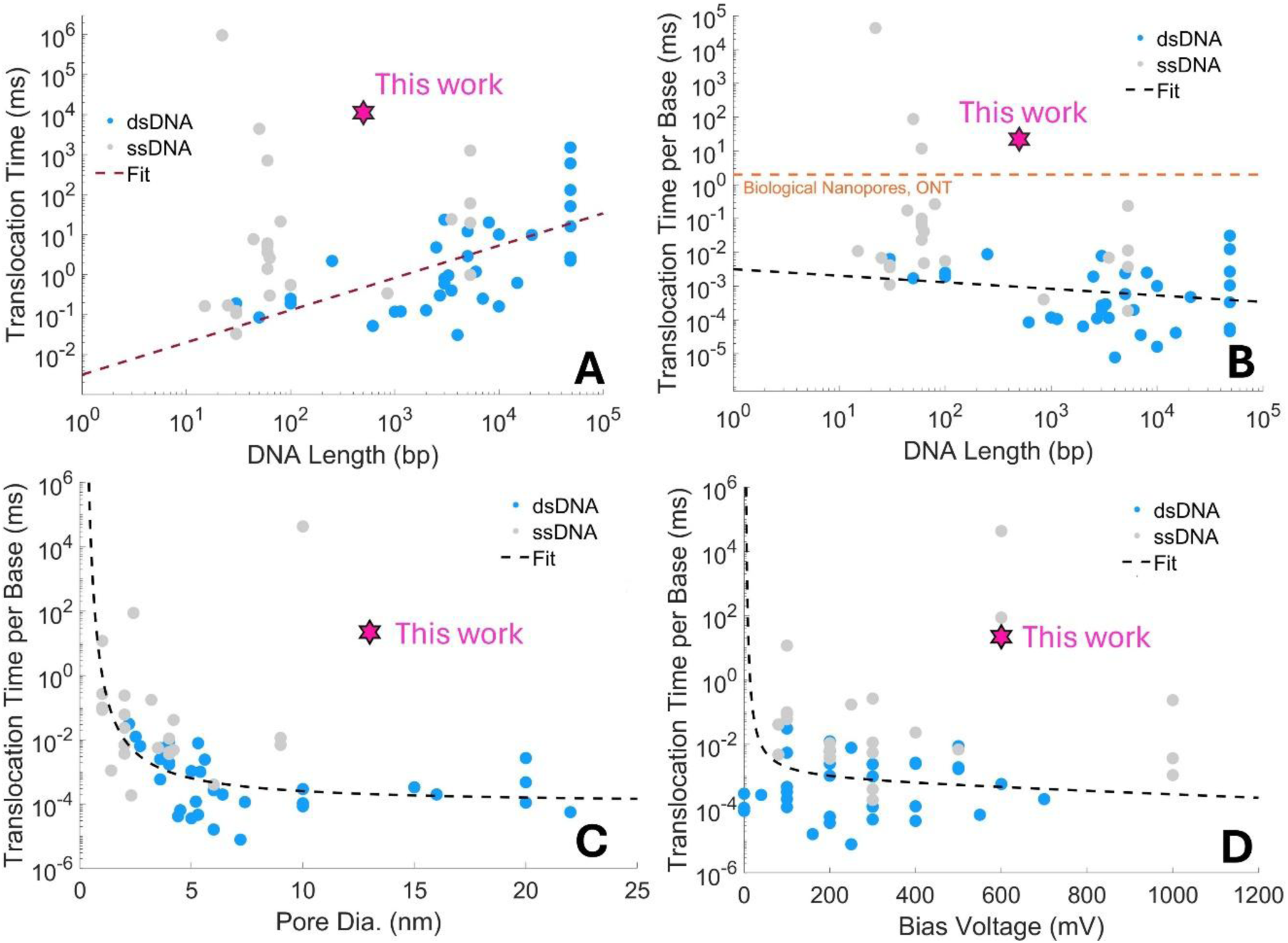
Comparison of the dwell-time reported in this work against other reports in literature. This work represents a substantial improvement in slowing down DNA translocation, specifically dsDNA, through solid-state nanopores achieving even slower slowdown compared to enzyme driven DNA translocation in biological pores. The references used to compile these plots are shown in Table S3 below.

#### Effect of DNA length

Figure S8A shows the reported dwell time as a function of DNA length from previous reports. The blue data points are from reports using double-stranded DNA (dsDNA) and the grey ones are from reports using single-stranded DNA (ss-DNA). We see an increase in the dwell-time with increasing DNA length as expected although on the short DNA length regime, dominated by reports using ssDNA, this correlation is weaker. We also note that some of the longest reported DNA translocation times are for short ssDNA. The dashed line in the figure shows a heuristic fit across the reported values. The function used for fitting is,

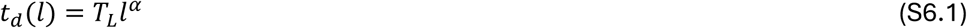

Where *t_d_* is the dwell time as a function of the DNA length *l* and *T_L_* and *α* are the fitting parameters. *T_L_* can be interpreted as the translocation time with *l* = 1, i.e. for a single DNA base. The parameters *T_L_* and *α* were obtained as 3.2 *μ*s and 0.35 respectively. The value of 3.2 *μ*s for *T_L_* is close to the typical DNA translocation speed of ∼ 1 – 10 *μ*s per base reported for DNA translocation in solid-state nanopores [27].

We also looked at the reported translocation times per base by dividing the reported translocation time with the length of the DNA used in the literature report. This is shown in Figure S8B. The fit function will now be *t_d_*(*l*) = *T_L_l^α^*^−1^. The translocation time per base allows us to compare these values with biological nanopore translocation rates reported for Oxford Nanopore Sequencers. These systems use an enzyme that unwinds and processes DNA through the pore at the rate of around 500 bases per second yielding a per based translocation time of 2 ms per base. From Figures 8A and 8B, we can note that this work represents a significantly reduced translocation speed for dsDNA compared to the state of the art and is even slower than that reported for biological pores (Figure 8B). There are 5 references [13, 15, 23, 26, 28 in Table S3] that report a translocation time greater than 200 ms, which was the threshold used in this work to classify “flat-like” vs long events. Out of these 5, all except ref. 23 use ssDNA, and all except 15 achieve slow translocation by using reduced pore diameter (1-2 nm) [See also SI Figure 8C]. It is interesting to note that reference 15 in Table S3 demonstrates slow down of short ssDNA (22 nt long) using electric field modulation around the pore, in a similar spirit as to our work.

#### Effect of pore diameter

Reducing the pore diameter can increase DNA-pore interactions and result in slower translocation. Indeed, when the dwell-times reported in literature are plotted against the pore diameter used in these experiments, we do see a reduction in translocation speed (or increase in dwell-time per base) with reduction in pore diameter. We use a heuristic fit function,

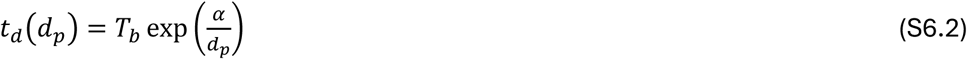

Where, *t_d_*(*d_p_*) is the dwell-time per base as a function of pore diameter *d_p_* and *α* and *T_b_* are fit parameters. This function is motivated by the expectation that *t_d_* should diverge (become infinity) as *d_p_* approaches 0 and reach a limiting value as *d_p_* becomes larger. Fitting Eq. S6.2 to the experimental data reported in literature yields *α* = 4 and *T_b_* = 0.2 *μ*s/base. Looking at Figure S8C, we can note that the present work represents a significant reduction in the translocation speed compared to the state of the art. Additionally, for the pore diameter used, the heuristic fit yields a dwell time of 0.15 ms for 500 bp dsDNA which closely matches the mean dwell time of 0.2 ms obtained for the fast, or “flat-like” events observed in our experiments.

#### Effect of bias-voltage

Figure S8D shows the comparison of dwell-times reported in literature as a function of bias voltage used. We do not see a strong effect of the bias voltage on the dwell times reported. However, motivated by the fact that the dwell time should diverge as the bias voltage, *V* approaches 0, and that a reduction is expected with increasing *V* due to increased electrophoretic force, we use the following function to get a heuristic fit to the experimental data,

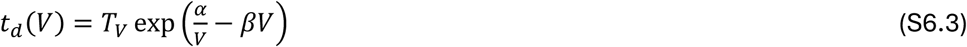

From the fit to the experimental values, *T_L_* is obtained as 0.8 *μ*s/base, close to what was obtained with Eqs. (S6.1) and (S6.2); *α* = 44 mV and *β* = 5×10^-4^ per mV.

**Supplementary Table S3.**
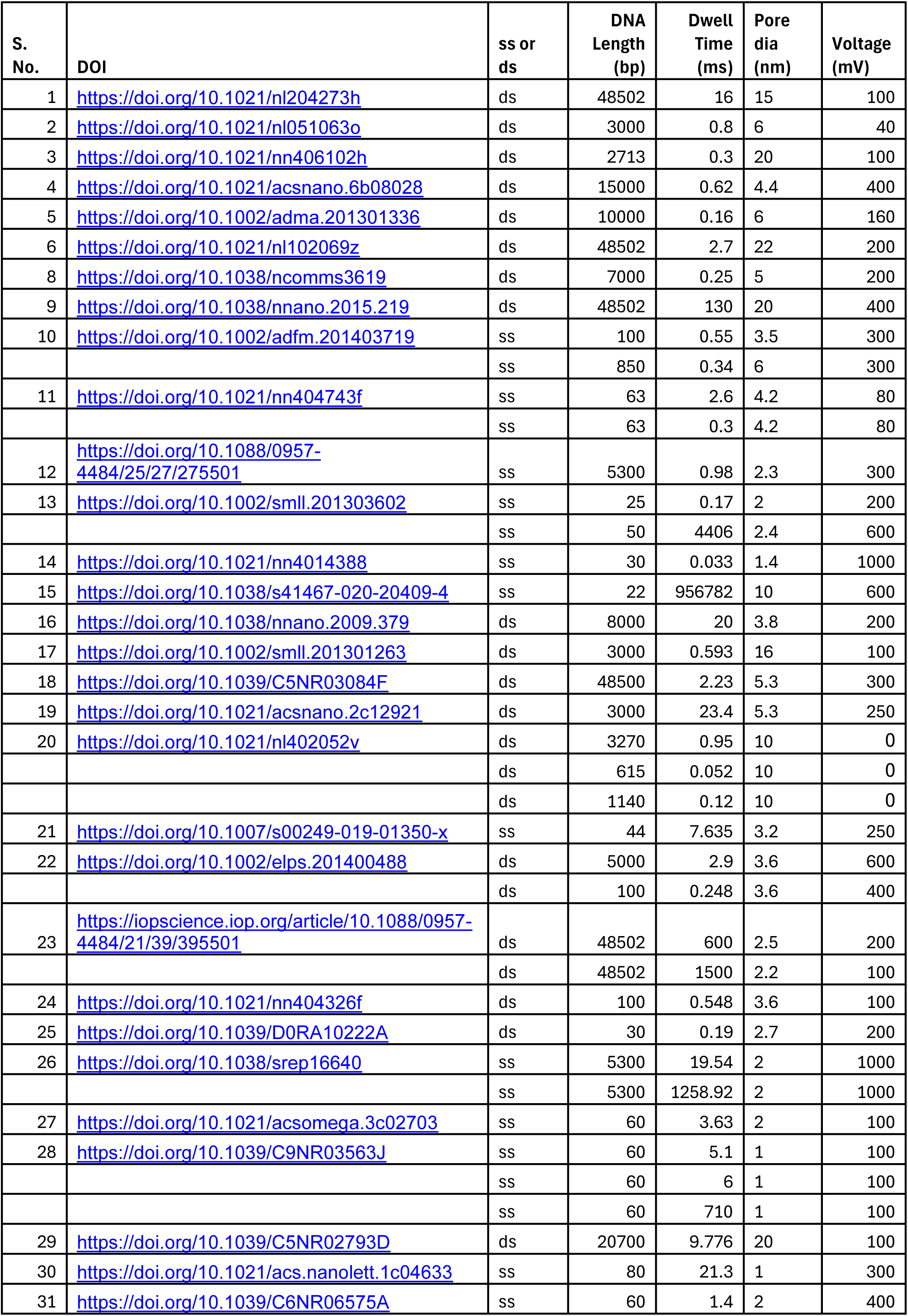
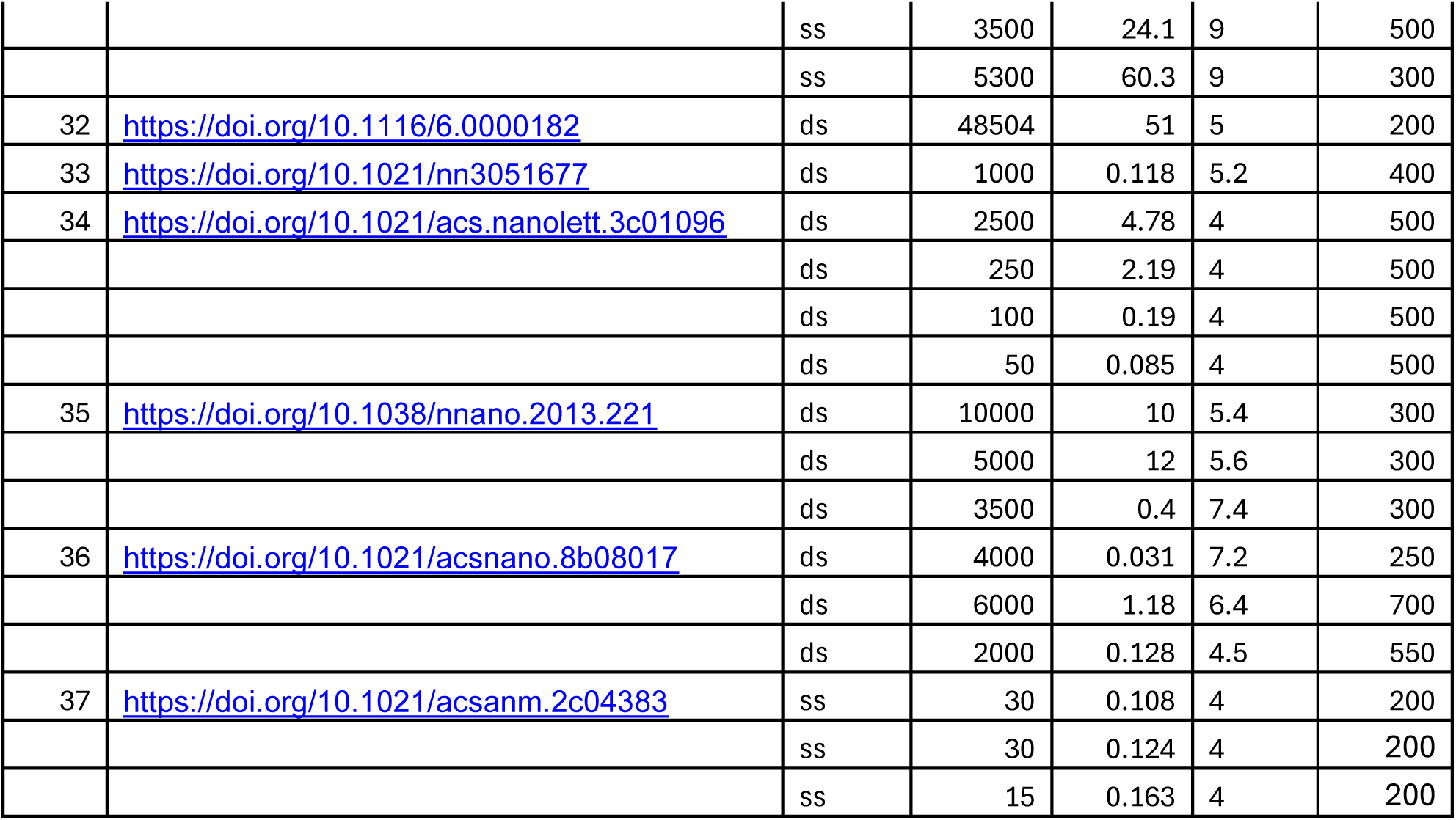
List of references used for benchmarking DNA translocation time reported in this work against other reports.

